# Shared roles of dorsal and subgenual anterior cingulate cortices in economic decisions

**DOI:** 10.1101/074484

**Authors:** Habiba Azab, Benjamin Y. Hayden

## Abstract

Theories of dorsal anterior cingulate cortex (dACC) function generally emphasize its cognitive and regulatory functions while theories of subgenual ACC (sgACC) emphasize its emotional, limbic, and arousal-related roles. But how different are these areas when compared in the same task? We recorded neuronal responses in both regions in macaques in a task with cognitive and limbic aspects, a token gambling task. Using tokens allowed us to compare responses to wins and losses. Both regions phasically encoded several important economic variables in similar ways; these included offered values, remembered values, attended values, and obtained values, and number of current tokens. Signal-to-noise ratio in sgACC was substantially lower than in dACC, and sgACC neurons responded more strongly to losses and in anticipation of large rewards. These results highlight the common economic functions of the anterior cingulum and suggest different functional emphases between regions, rather than a strict cognitive vs. emotional division.

## INTRODUCTION

The cingulum is a large structure on the medial wall encircling the corpus callosum in the saggital plane. The portion rostral to the central sulcus, the anterior cingulate cortex (ACC), is important for almost every aspect of cognition and its malfunction is associated with several psychiatric diseases (Bush et al., 1999; Heilbronner & Hayden, 2016; Paus, 2001; Rushworth et al., 2012; Shenhav et al., 2013). A great deal of evidence indicates that the ACC can be functionally subdivided into at least two portions, a dorsal (dACC) and subgenual (sgACC) one. These portions are associated with different cytoarchitectonics, connectivity patterns, patterns of hemodynamic activation, and lesion effects (Bush et al., 2000; Vogt et al., 1995). Nonetheless, a direct comparison of their function at the single-neuron level has never been made.

Theories of dACC generally emphasize its cognitive and regulatory roles rather than limbic and emotional ones. Cognitive functions associated with dACC include error- and conflict-monitoring, allocation of control, task-switching and regulation of strategy (Allman et al., 2001; Amiez et al., 2005; Carter et al., 2007; Kerns et al., 2004; Neubert et al., 2015; Procyk et al., 2000; Quilodran et al., 2008; Rushworth et al., 2002; Shenhav et al., 2013). In contrast, theories of sgACC emphasize its limbic, emotional, and arousal-related roles and not its cognitive function. These functions include emotional processing, mood-changes, sleep, and potentially a role in depression (Botteron et al., 2002; Coryell et al., 2005; Drevets et al., 1997; George et al., 2006; Johansen-Berg et al., 2008; Rolls et al., 2003). Some evidence suggests that these regions have negatively correlated activity patterns (Bush et al., 2000), consistent with the idea they inhibit each other. These differences have led to the theory that these two regions play antagonistic roles in cognition (Bush et al., 2000).

Nonetheless, there are reasons to believe that dACC and sgACC play overlapping roles in cognition. Both are associated with monitoring and anticipation of reward (which itself has both cognitive and limbic aspects.) The role of the dorsal portion of ACC in reward representation is well-established (Heilbronner and Hayden, 2016). Although single-unit studies in the subgenual portion have been relatively few, some of these studies also implicate this region in reward processing and representation (Monosov & Hikosaka, 2012; Rudebeck et al., 2014; similar findings were reported in Amemori & Graybiel, 2012, who recorded in the adjacent pregenual cingulate). More broadly, both regions have been implicated in pain processing (Derbyshire et al., 1998; Vogt et al., 1992), depression (Cotter et al., 2001; Mayberg et al., 1997), addiction (Forman et al., 2004; Goldstein et al., 2007), OCD (Graybiel & Rauch, 2000), and other psychiatric disorders (Blumberg et al., 2000; Bouras et al., 2001; Drevets et al., 1997; Drevets et al., 2008; Ongur, Drevets & Price, 1998). Nonetheless, our ability to compare these regions is limited by the paucity of direct comparisons of their function.

Here we compared the responses of these two regions in a single gambling task. We sought to understand their relative contributions within a single task that was complex and engaging, and had both limbic and cognitive aspects. Among economic choice tasks, gambling is particularly useful because it is well-characterized mathematically, preferences are stable, and it involves both reward and monitoring/adjustment processes (Heilbronner and Hayden, 2013; Heilbronner & Hayden, 2015). Furthermore, we have a foundational understanding of the neuroscience of risky choice, including recordings in dACC, although not in sgACC (Blanchard and Hayden, 2014; Hayden et al., 2011; Hosokawa et al., 2013; Kennerley et al., 2009; Procyk et al., 2000). To directly compare coding of gain and loss, we added a new feature to our task. On each trial, monkeys gambled for gain or loss of tokens; collection of 6 tokens produced a jackpot reward (cf. Seo & Lee, 2007). This feature allowed for direct comparison of two reward signs in a single modality (thus controlling for sensory features that, say, an air puff or hypertonic solution would not). This manipulation also allowed us to explore responses to secondary reinforcers—which has not been done in sgACC.

These two cingulate regions exhibited strikingly similar patterns of activity in our study. Both encoded several economic variables and did so in similar ways. Specifically, we found encoding of the values of offers (attended and remembered), outcomes, and number of tokens accumulated. Both regions used an attentionally-aligned, as opposed to labeled line, coding scheme for values. We also confirmed earlier results showing encoding of positions of offered and chosen options (Strait et al., 2016). We found no measurable differences in response latency. This is not to say the regions were identical: two major differences stood out. First, task-relevant coding was more frequent in dACC and signal-to-noise was roughly three times greater. Second, neurons in dACC were biased towards higher firing for gamble wins, while neurons in sgACC were biased towards higher firing for losses. There was also a positive bias at the single neuron level in anticipation of large, primary rewards in sgACC; no corresponding bias occurred in dACC. These data suggest that dACC and sgACC play overlapping roles in economic decisions albeit with different emphases, but do not have antagonistic roles or strong cognitive / emotional specialization.

## RESULTS

### Behavior

Two monkeys performed a token-gambling task. On each trial, monkeys chose between two options presented asynchronously that differed in three dimensions: win amount, loss amount, and probability of win (Figure 1A). Both monkeys showed behavior consistent with task understanding (Figure 2). Specifically, both subjects preferred the option with the greater expected value (subject B: 80.3%; subject J: 75.1%; both P < 0.0001; two-sided binomial test), and were sensitive to the three parameters that defined each gamble (See Table 1). Subjects were more sensitive to win amount than to loss amount, consistent with the idea that they focus on the win (Hayden & Platt, 2007; Hayden et al., 2008). Monkeys showed weak side and order biases (rightward choice: 51.0% in subject B; 47.4% in subject J; chose second: 52.5% in subject B; 55.6% in subject J.)

**Figure 1:**
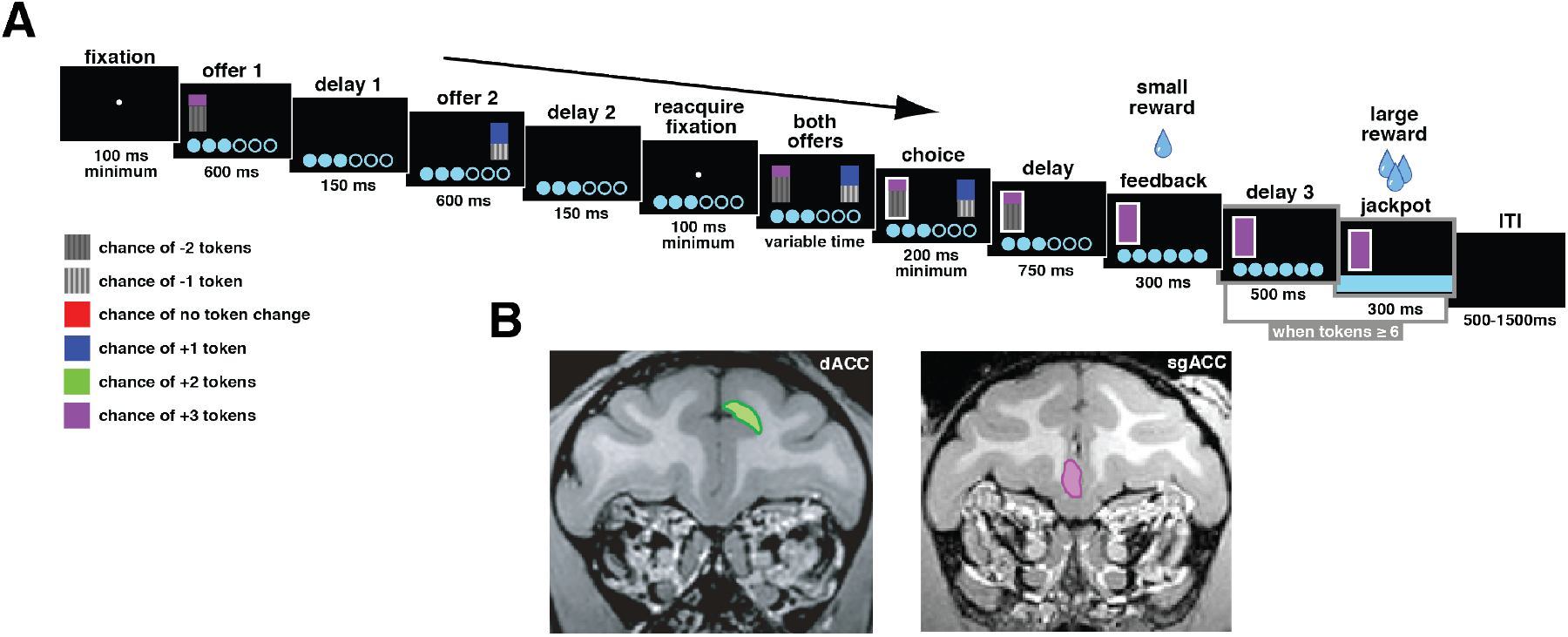
**A:** Example trial from token-gambling task. Offers were presented asynchronously, and the side the first offer appeared on was counterbalanced across trials. Each gamble offered one of two outcomes (indicated by the colors of the bars) at a certain probability (indicated by their heights). **B:** Regions of interest (for exact coordinates, see Methods).

**Figure 2:**
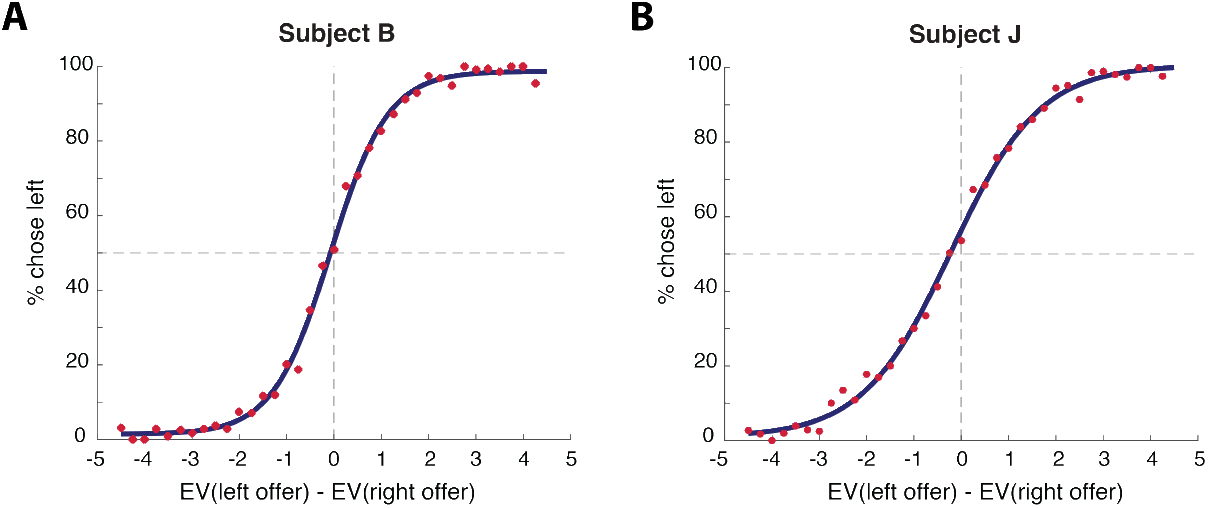
Behavior for each subject, fit to a sigmoid function. Subjects choose the left option more often as its value increases. EV: expected value of gamble.

**Table 1:** Regression coefficients from a logistic regression model of choice (left = 1, right = 0) against the variables in the regressor column. A ‘loss’ within a gamble was always less than or equal to the win outcome of that gamble. Note that losses included negative outcomes. Overall, the expected value of the win-portion of a gamble influenced behavior more than the value of a loss did in both subjects.

| Regressor | Subject B | Subject J |
| --- | --- | --- |
| Win amount, left offer | 1.2302 (P < 0.0001) | 1.0097 (P < 0.0001) |
| Loss amount, left offer | 0.4063 (P < 0.0001) | 0.2451 (P < 0.0001) |
| Probability of win, left offer | 3.3828 (P < 0.0001) | 3.1745 (P < 0.0001) |
| Win amount, right offer | -1.1577 (P < 0.0001) | -1.1390 (P < 0.0001) |
| Loss amount, right offer | -0.3449 (P < 0.0001) | -0.1937 (P < 0.0001) |
| Probability of loss, right-offer | -4.1934 (P < 0.0001) | -3.5026 (P < 0.0001) |
| Intercept | -0.2893 (P < 0.0001) | 0.6177 (P < 0.0001) |

Monkeys were risk-seeking, consistent with earlier studies (Hayden et al., 2008; Hayden & Platt, 2007; McCoy & Platt, 2005). To assess risk-seeking, we fit a utility curve distortion parameter to subjects’ behavior in each session (Lak et al., 2014; Yamada et al., 2013). This parameter, α, was always greater than 1 in both subjects (B: α = 1.61 (n = 66 sessions); subject J: 1.81 (n = 74 sessions); both greater than 1 at P < 0.0001; one-sample t-test over sessions). Subjects’ behavior was modestly influenced by the number of current tokens: accuracy improved as tokens increased (subject B: β = 0.027, P = 0.0012; subject J: β = 0.078, P < 0.0001; logistic regression of accuracy against tokens acquired).

### Variance explained by task variables is higher in dACC than in sgACC

Responses to task variables were attenuated in sgACC neurons compared to dACC neurons. To quantify this difference, we regressed average firing rate in a sliding 500 ms window against eight task-relevant variables (see Methods). We then computed the average fraction of variance in firing rates explained by this model for each neuron in each brain region over the entire trial. We next compared these measures across regions, where neurons were the unit of analysis. Task-relevant variables explained over three times as much variance in dACC than in sgACC neurons (average adjusted R^2^ over all neurons: dACC: 0.0093; sgACC: 0.0025; T = 4.94, P < 0.0001; two-sample t-test). Thus, variance explained was between 3 and 4 times greater in dACC than in sgACC.

### Neurons in both regions encode offered values

Responses of two neurons (one in dACC and one in sgACC) are shown in Figure 3. Both of these neurons’ firing rates were correlated with the expected value of the first offer while it was displayed on the screen (epoch 1). The firing rate of the dACC neuron was also affected by the expected value of the first offer when the second offer appeared (epoch 2). This was not the case for the sgACC neuron.

**Figure 3:**
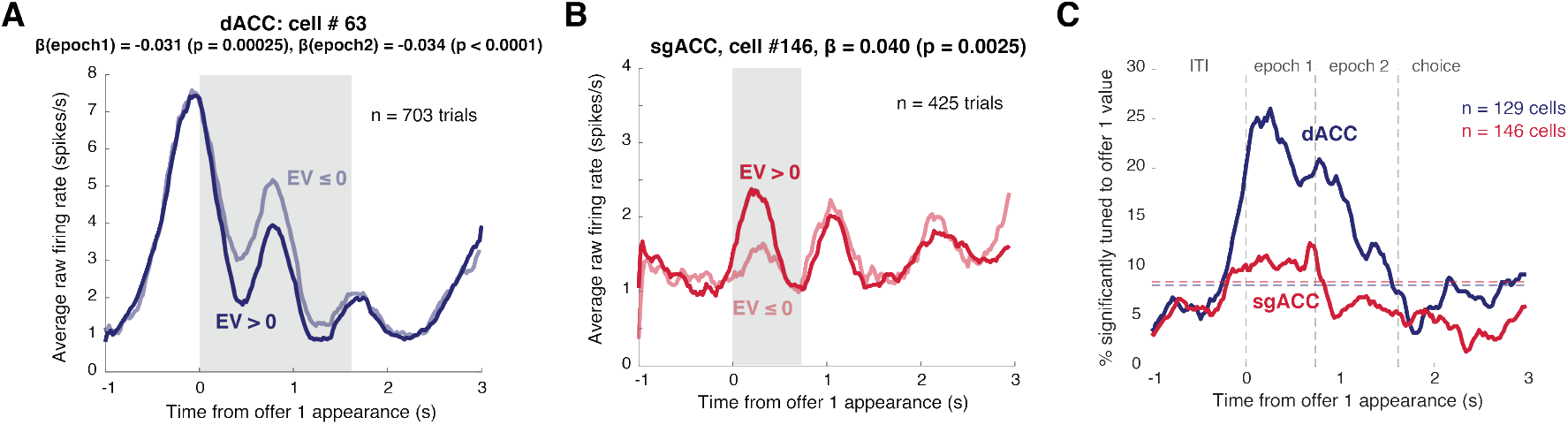
Neural responses to the value of offer 1. **A:** Responses of one dACC neuron with increasing firing rate in response to smaller values of offer 1 in both the first and second epochs. Shaded regions are epochs of interest with significant modulation. **B:** Responses of one sgACC neuron with increasing firing rates in response to larger values of offer 1 in the first epoch. **C:** Proportion of neurons selective for the value of offer 1 in both regions throughout the course of the trial. This fraction was computed by regressing firing rate over a 500 ms sliding window against the expected value of the first offer, along with other task-relevant variables (see Methods). The blue and red horizontal lines indicate the percentage of neurons we would expect to see by chance (as determined by a two-sided binomial test, according to the number of neurons in each area) for dACC and sgACC, respectively.

Significant proportions of neurons in both regions encoded the value of the first offer in the 500 ms epoch starting 100 ms after that offer appeared. Note that we chose this epoch for another study and use it here to allow comparability across datasets (Strait et al., 2014; Strait et al., 2015).

We found that responses of 25.6% (n = 33/129) of neurons in dACC and 12.3% (n = 18/146) in sgACC were affected by the value of offer 1 during epoch 1. Both proportions are greater than chance (dACC: P < 0.0001; sgACC: P = 0.0004; two-sided binomial test). Figure 3C shows the fraction of neurons whose firing rates were modulated by offer 1 value through time. There was no measured bias towards a positive direction in either area (see Table 3). The average regression weight across all neurons, another measure of directionality of effect, was not different from zero in either area, indicating no bias in the overall populations (see Table 2).

**Table 2:** Assessment of biases in tuning in the overall population. We performed this analysis running a one-sample t-test on the regression coefficients corresponding to each variable obtained from a multiple linear regression (see Methods).

| Variable | dACC (t-statistic) | sgACC (t-statistic) |
| --- | --- | --- |
| Offer 1 value (attended) | 0.64 (P = 0.53) | 1.67 (P = 0.098) |
| Offer 1 value (remembered) | -0.26 (P = 0.79) | -0.47 (P = 0.64) |
| Offer 2 value (attended) | 0.95 (P = 0.34) | 0.97 (P = 0.33) |
| Chosen value | -1.47 (P = 0.14) | **-2.31 (P = 0.022)** |
| Unchosen value | **2.70 (P = 0.0078)** | 1.73 (P = 0.085) |
| Outcome | **2.35 (P = 0.02)** | **-2.85 (P = 0.0050)** |
| Number of tokens | 1.42 (P = 0.16) | 0.15 (P = 0.88) |
| Jackpot | 0.092 (P = 0.93) | **4.47 (P < 0.0001)** |

**Table 3:** Assessment of biases in frequency of coding in the tuned population. We determine whether the fraction of positively tuned neurons (of the tuned population only) is significantly larger than chance (50%), using a two-sided binomial test.

| Variable | dACC (% positive) | sgACC (% positive) |
| --- | --- | --- |
| Offer 1 value (attended) | 57.6% (n = 19/33, P = 0.49) | 66.7% (n = 12/18, P = 0.24) |
| Offer 1 value (remembered) | 56.0% (n = 14/25, P = 0.69) | 41.7% (n = 5/12, P = 0.77) |
| Offer 2 value (attended) | 50.0% (n = 9/18, P = 1) | 57.1% (n = 8/14, P = 0.79) |
| Chosen value | **20% (n = 4/20, P = 0.012)** | 22.2% (n = 2/9, P = 0.18) |
| Unchosen value | **100% (n = 12/12, P = 0.0005)** | 50.0% (n = 3/6, P = 1.0) |
| Outcome | **71.0% (n = 22/36, P = 0.029)** | 31.6% (n = 6/19, P = 0.17) |
| Number of tokens | 60.0% (n = 27/49, P = 0.23) | 58.3% (n = 14/31, P = 0.54) |
| Jackpot | 54.3% (n = 19/35, P = 0.74) | 70.6% (n = 12/17, P = 0.14) |

### Neurons in both regions integrate gamble probability and stakes

The mathematical expected value of a gamble in our task – and its subjective value – depend on three factors: the probability of the win, the win amount, and the loss amount. We wanted to know whether the individual components of a gamble are integrated into a single general value signal, or represented as separate attributes.

We explored whether probability and stakes were integrated in the first epoch of the trial, when only the first gamble had been displayed. We compared regression weights for neuronal responses to the first offer against the magnitude and probability of a win and the magnitude of loss, in one multivariate regression, while controlling for other variables that significantly explained firing rate for this neuron in this epoch (see Methods for further detail).

Regression coefficients corresponding to win-magnitude and win-probability were positively correlated, supporting the hypothesis that these parameters are integrated This correlation is observed in both cingulate regions (dACC: r = 0.63, P < 0.0001; sgACC: r = 0.31, P = 0.0001; Pearson correlation coefficient of signed regression coefficients). The absolute values of these coefficients were also positively correlated, indicating that these variables were encoded by a single population of neurons rather than two distinct ones — further supporting the integration hypothesis (dACC: r = 0.59, P < 0.0001; sgACC: r = 0.56, P < 0.0001). We previously reported similar findings in the ventromedial prefrontal cortex (vmPFC) and the ventral striatum (VS, Strait et al., 2015). These results suggest that all four regions carry integrated value signals.

Win and loss magnitudes were also integrated in both areas (dACC: r = 0.34, P = 0.0001; sgACC: r = 0.36, P < 0.0001; Pearson correlation coefficient of signed regression coefficients). While these variables were represented in overlapping populations in sgACC (r = 0.30, P = 0.0003; Pearson correlation coefficient of unsigned regression coefficients), this relationship does not achieve significance in dACC (r = 0.17, P = 0.053).

### Neurons in both regions encode remembered values

We examined whether firing rates reflected the value of the first offer while the second offer was presented (epoch 2, 500 ms epoch starting 100 ms after the second offer appeared,). We saw encoding of offer 1 in this epoch in dACC (19.4% of neurons, n = 25/129, P < 0.0001; two-sided binomial test), but it failed to reach significance in sgACC (8.22%, n = 12/146, P = 0.084). Since task-relevant signals were generally attenuated in sgACC, and the percentage of tuned neurons we saw failed to reach significance by one neuron, we repeated this analysis in an earlier 500 ms epoch, starting immediately when the second offer appeared. The fraction of neurons tuned to the value of offer 1 in this epoch does achieve significance (11.0%, n = 16/146, P = 0.0028; two-sided binomial test). This proportion is still significant when correcting for multiple comparisons (i.e. at P < 0.025). This finding suggests that sgACC may, in fact, carry a memory signal, although one that is more attenuated and less reliable than that in dACC. There was no bias in tuning in either brain region (see Table 3), nor was there any bias in the regression coefficients of the overall populations (see Table 2).

We next investigated the relationship between the pattern of tuning for attended and remembered offers. Was offer 1 encoded in similar tuning formats as it moved from the retina to working memory? And were the putative working memory neurons, which carried the value of offer 1 in the second epoch, the same ones that encoded it when it was first presented—in the first epoch? The answers to both questions is yes for dACC, but we cannot draw conclusions from our results in sgACC. We computed regression weights for the expected value of offer 1 in epoch 1 (when it was present on the screen) and for the same option in epoch 2 (when it was remembered), while controlling for other task variables that best explained each neuron’s response (see Methods). These values were positively correlated in dACC (r = 0.32, P = 0.0002; Pearson correlation coefficient of signed regression coefficients), indicating a preservation of coding format as information was transferred from an active to a remembered storage buffer. The absolute regression coefficients were also positively correlated (r = 0.44, P < 0.0001), suggesting these variables were encoded by a single task-sensitive population, rather than memory and active-buffer neurons (consistent with vmPFC and VS; Strait et al., 2015).

We cannot draw any strong conclusions from results in sgACC. There was no significant correlation between these variables: neither one indicating preservation of format (r = 0.090, P = 0.28; Pearson correlation of signed regression coefficients), nor one indicating that this signal was carried by the same population (r = 0.0380, P = 0.65; Pearson correlation of unsigned regression coefficients). Results were similar in the slightly earlier 500 ms epoch we used earlier (preservation of format: r = 0.15, P = 0.079; overlapping populations: r = 0.063, P = 0.45). It is difficult to draw conclusions from the absence of this correlation, as it could be due to weaker signal-to-noise ratio in this region. We therefore performed a cross-validation procedure similar to that in Blanchard et al. (2015), and found that our signal in the epochs of interest was not strong enough to detect correlations (see Methods). The failure of this procedure suggests that we could not detect this effect with our data, even if it existed. We therefore draw no conclusions from this null result.

### Simultaneous encoding of both offer values

We next explored the effects of the second offer value on firing rates during epoch 2. Figure 4 shows two example neurons, one from dACC and one from sgACC, whose firing rates increased in response to the second offer. A significant proportion of neurons in both regions encoded the value of the second offer when it appeared (dACC: 14.0%, n = 18/129; sgACC: 9.59%, n = 14/146). Both proportions exceed what we would predict by chance (dACC: P = 0.0001; sgACC: P = 0.016; two-sided binomial test). There was no significant bias in tuning in either area (Table 3), and the average coefficient of offer 2 value over the entire population was also not biased in either brain region (Table 2). Figure 4C shows how the fraction of neurons responsive to offer 2 changed over the course of the trial.

**Figure 4:**
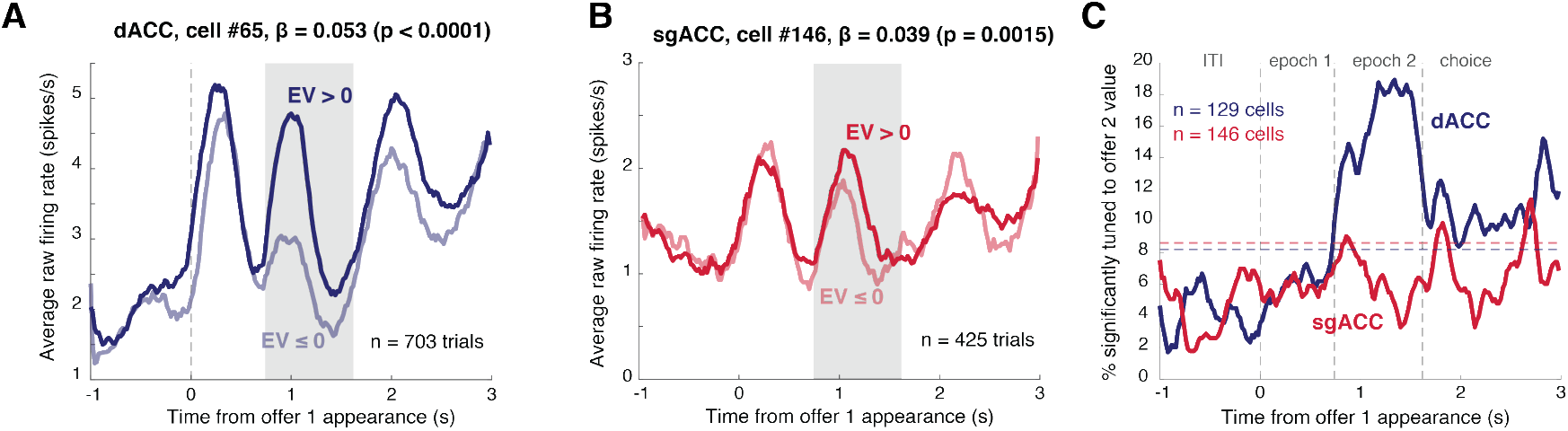
Neural responses to the value of offer 2. **A:** Responses of one dACC neuron whose firing rate increased in response to larger values of offer 2. **B:** Responses of one sgACC neuron whose firing rate increased in response to larger values of offer 2. **C:** Percentage of neurons tuned to the value of the second offer in both regions over the course of the trial. A 500 ms sliding window was used to compute these values at each point of the trial. The blue and red horizontal lines indicate the percentage of neurons we would expect to see tuned by chance (as determined by a two-sided binomial test, according to the number of neurons in each area) for dACC and sgACC, respectively.

### Attentionally-aligned coding scheme for offer values

Do cingulate regions encode all attended offers similarly, using the same population? Our previous studies of vmPFC and VS indicate that value coding in these regions is attentionally-aligned – that is, they encode the value of the attended offer in a conserved attended format (by format, we mean tuning strength and sign, and identity of neurons; Strait et al., 2015; Akaishi & Hayden, 2016; Rich & Wallis, 2016).). This framework differs from a labeled-line format, in which a specialized set of neurons encodes the value of each offer respectively (e.g. Hunt et al., 2015). To test this, we compared the regression coefficients associated with the first offer in the first epoch to those associated with the second offer in the second epoch.

Our data support the attentional alignment hypothesis. Specifically, coding direction for both offers was consistent in both areas (dACC: r = 0.51, P < 0.0001; sgACC: r = 0.26, P = 0.0013; Pearson correlation test of signed coefficients). Second, both regions made use of a consistent set of neurons for the two offers (dACC: r = 0.45, P < 0.0001; sgACC: r = 0.32, P = 0.0001; Pearson correlation test of unsigned coefficients).

### Beta anti-correlation in dACC: a putative signature of comparison through mutual inhibition

One putative neural signature of the comparative process is an anti-correlation between regression coefficients for the values of the two offers while they are being compared (Strait et al., 2014). This anti-correlation indicates that the two offers modulate neuronal activity in opposing directions, and that the ensemble of neurons effectively subtracts their values. We observed this pattern in dACC during the second epoch—when both offers had been presented (r = −0.17, P = 0.050; Pearson correlation coefficient of signed regression coefficients), suggesting that this area is involved in comparing the two offers. We found no significant correlation in sgACC (r = −0.077, P = 0.35). It is difficult to interpret the lack of a significant correlation as the absence of a comparison signal in light of our weak signal-to-noise ratio, and due to the failure of the cross-validation procedure (see Methods).

### Encoding of chosen and unchosen value signals in both regions

We examined selectivity for chosen and unchosen offer values in the 1-second epoch ending with the saccade that indicated the subject’s choice. We chose this epoch to capture any correlates of the choice process that arose when the second offer was presented (presumably when the comparison was made), and when the subject had to indicate his choice.

Figure 5 shows four neurons whose firing rates were affected by the values of the chosen and unchosen offer values in both brain regions. Significant proportions of such neurons were found in dACC, but not in sgACC. 15.5% (n = 20/129) of dACC responded to the chosen value (two-sided binomial test: P < 0.0001), and 9.30% (n = 12/129) responded to the unchosen value (P = 0.030). The proportion of neurons encoding chosen value was greater than the proportion encoding unchosen value (P = 0.018; two-sided binomial test). However, encoding of chosen value was not stronger than encoding of unchosen value when the tuning coefficients of all neurons in the population were compared (T = 1.67, P = 0.097; unpaired two-sample t-test of absolute regression coefficients). There was more overlap than expected by chance between these populations (r = 0.22, P = 0.013; Pearson correlation of unsigned regression coefficients). Figure 5E shows the change in these fractions as the choice was made.

**Figure 5:**
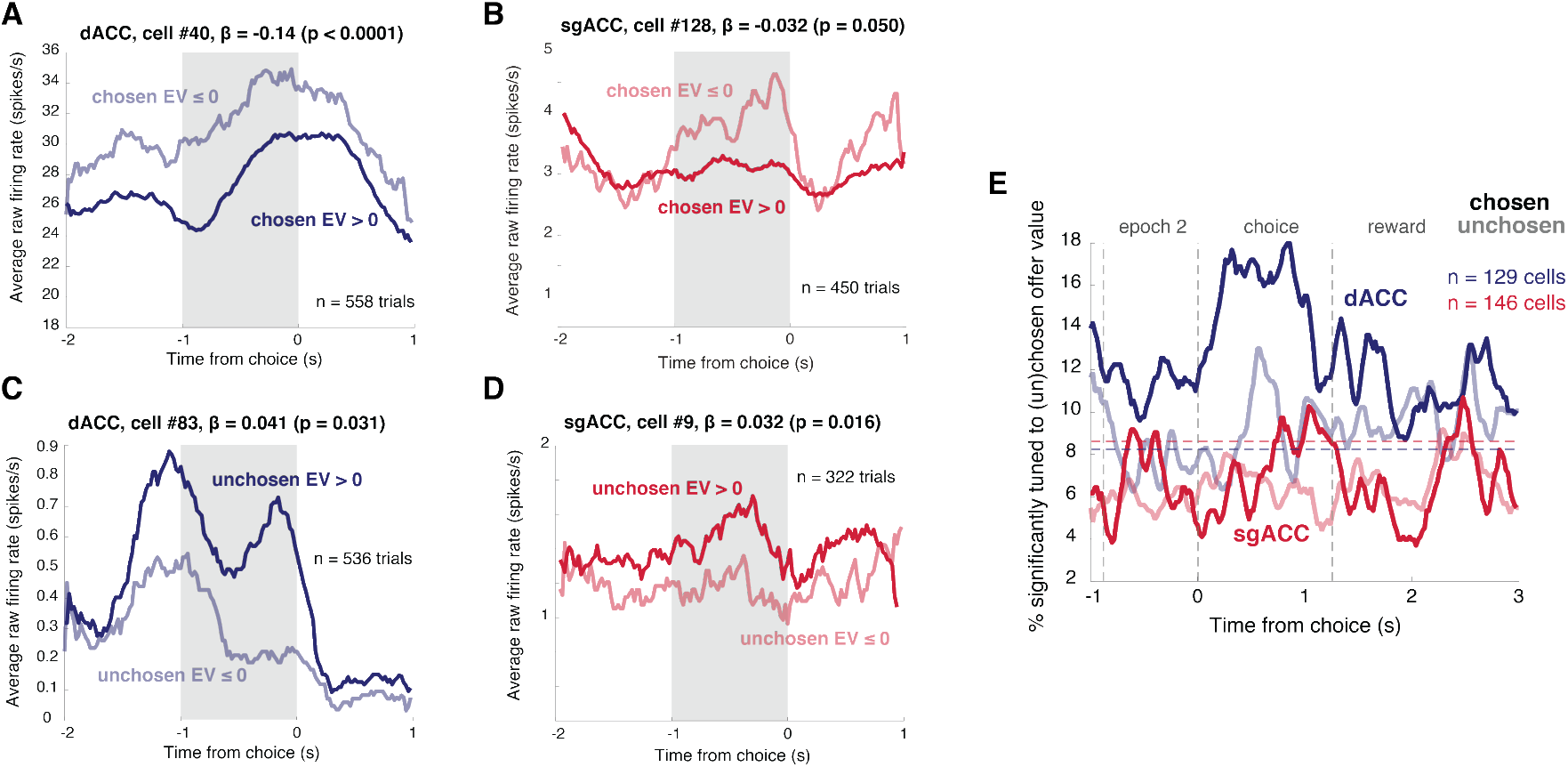
Neural responses to the values of the chosen and unchosen offers. **A:** Responses of dACC neuron whose firing rate was negatively modulated by the value of the chosen offer. This pattern is typical of neurons in this region (see text and Tables 2 and 3). **B:** Responses of sgACC neuron whose firing rate was negatively modulated by the value of the chosen offer. **C:** Responses of dACC neuron whose firing rate was positively modulated by the value of the unchosen offer. This pattern is typical of neurons in this region (see text and Tables 2 and 3). **D:** Responses of sgACC neuron whose firing rate was positively modulated by the value of the unchosen offer. **E:** Percentage of neurons tuned to the values of the chosen and unchosen offer values in both cingulate regions throughout the course of the trial. Percentages were computed using a 500 ms sliding window, using a regression analysis (see Methods for details). The blue and red horizontal lines indicate the percentage of neurons we would expect to see tuned by chance (as determined by a two-sided binomial test, according to the number of neurons in each area) for dACC and sgACC, respectively.

There was a bias towards negative tuning for the chosen value in the tuned population (n = 16/20 negatively tuned, P = 0.012; two-sided binomial test), but not in the regression coefficients of the overall population (Table 2). We found a bias towards positive tuning for the unchosen value: both in the tuned population (n = 12/12 positively tuned; P = 0.0005; two-sided binomial test), and in the overall population (T = 2.70, P = 0.0078; one-sample t-test).

No significant proportion of neurons encoding chosen (6.16%, n = 9/146, P = 0.57; two-sided binomial test) or unchosen (4.11%, n = 6/146, P = 0.71) offer values was found in sgACC. However, the overall population was negatively biased towards encoding of the chosen value (T = −2.31, P = 0.022; one-sample t-test), and exhibited a positive trend in encoding the unchosen value (Table 2), echoing the patterns we see in dACC, albeit attenuated.

### Encoding of gamble outcomes

We examined selectivity for gamble outcomes during the 800 ms reward epoch that began with the reveal of the outcome and extended into the intertrial interval (see Figure 1). Recall that outcomes in this task were tokens, not juice rewards. Figure 6 shows responses of two example neurons, one from each region, selective for outcome amount. We excluded jackpot trials from this analysis so that liquid reward was matched in all cases; the only difference was tokens given or taken. 24.03% (n = 31/129) of neurons in dACC and 13.0% (n = 19/146) of neurons in sgACC encoded the outcome of a trial. Both of these proportions are significant (P < 0.0001 in dACC and P = 0.0001 in sgACC; two-sided binomial test). Figure 6C shows the peak of these fractions in the outcome epoch. These are the first demonstration of sgACC selectivity to gains or losses of secondary reinforcers.

**Figure 6:**
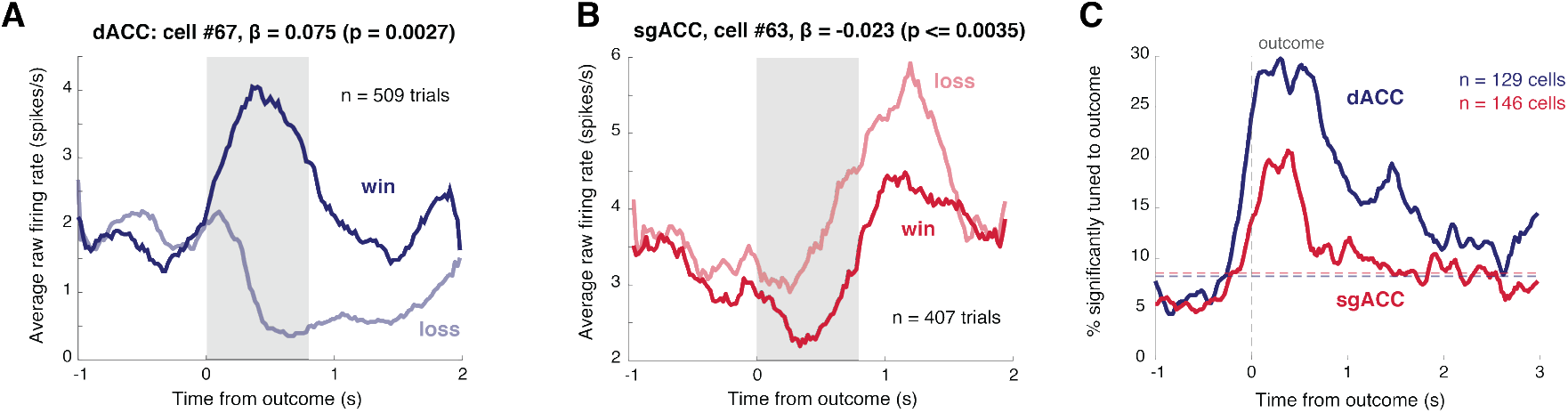
Neural responses to outcomes in both regions. **A:** Responses of dACC neuron with increasing firing rate in response to wins. There was a bias towards this pattern in the dACC population (see text and Table 2). **B:** sgACC neuron with increasing firing rate in response to losses. There was a bias towards this pattern in the sgACC population (see text and Table 2). **C:** Percentage of neurons in both regions tuned to the outcome of a trial throughout the course of a trial, using a 500 ms sliding window. The blue and red horizontal lines indicate the percentage of neurons we would expect to see tuned by chance (as determined by a two-sided binomial test, according to the number of neurons in each area) for dACC and sgACC, respectively.

Neurons in dACC showed a positive bias in outcome encoding. Specifically, the majority of tuned neurons were positively tuned for outcome (71.0% positively tuned, n = 22/36, P = 0.029; two-sided binomial test). There was also a positive bias in regression coefficients in the overall population (T = 2.35, P = 0.02; one-sample t-test). In sgACC, neurons in the overall population were negatively tuned to outcomes (T = −2.85, P = 0.0050; one-sample t-test), although this bias in the overall population’s regression weights was not significant in the tuned population (Table 3).

### Both regions encode current number of tokens

In our task, the number of tokens provided a measure of progress towards the jackpot reward. As noted, performance improved modestly with number of tokens. We hypothesized that this variable would affect neuronal responses in both cingulate areas. Figure 7 shows two neurons (one in dACC, the other in sgACC) whose responses were modulated by the number of tokens acquired during the 800 ms reward epoch. 34.9% (n = 45/129) in dACC and 16.4% in sgACC (n = 24/146) significantly represented this variable in that epoch. Both of these fractions are significant (P < 0.0001 in both regions; two-sided binomial test). Figure 7C shows the change in this fraction throughout the trial. There was no bias in tuning in either brain region: neither within the tuned population (Table 3), nor in the overall population’s regression coefficients (Table 2).

**Figure 7:**
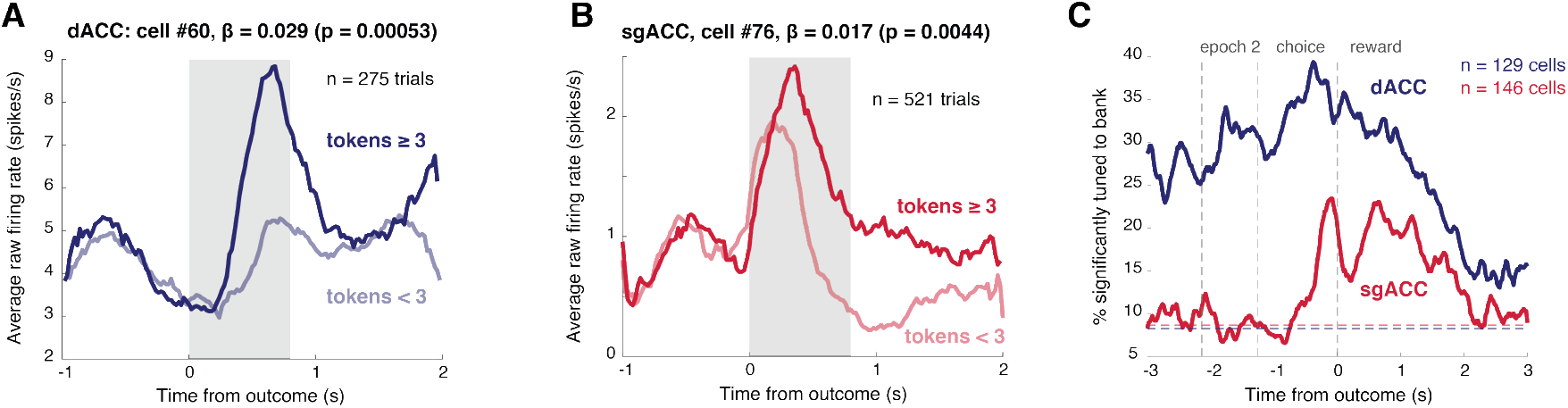
Neural responses to the number of tokens accumulated at the beginning of the trial. **A:** Responses of dACC neuron that increased its firing rate as subjects accumulated more tokens. **B:** Responses of sgACC neuron that increased its firing rate as subjects accumulated more tokens. **C:** Percentage of neurons in each area tuned to the number of tokens accumulated throughout the course of the trial. All percentages were computed using a 500 ms window (see Methods). The blue and red lines indicate the percentage of neurons we would expect to see tuned by chance (as determined by a two-sided binomial test, according to the number of neurons in each area) for dACC and sgACC, respectively.

### Neurons in both regions carry reward anticipation signals

We quantified neural responses to anticipation of jackpots by comparing them to responses on standard trials. Figure 8 shows two example neurons—one from each region—with activity that varied with anticipation of a jackpot reward. Significant proportions of such neurons were found in both regions (27.3% in dACC, n = 35/129, P < 0.0001; 11.6% in sgACC, n = 17/146, P = 0.0011; two-sided binomial test). We saw no bias in positive or negative tuning in dACC: neither in the tuned population (Table 3), nor in the population overall (Table 2). However, there was a positive bias in sgACC in the overall population (T = 4.47, P < 0.0001; one-sample t-test), although not significantly in the tuned population (Table 3).

**Figure 8:**
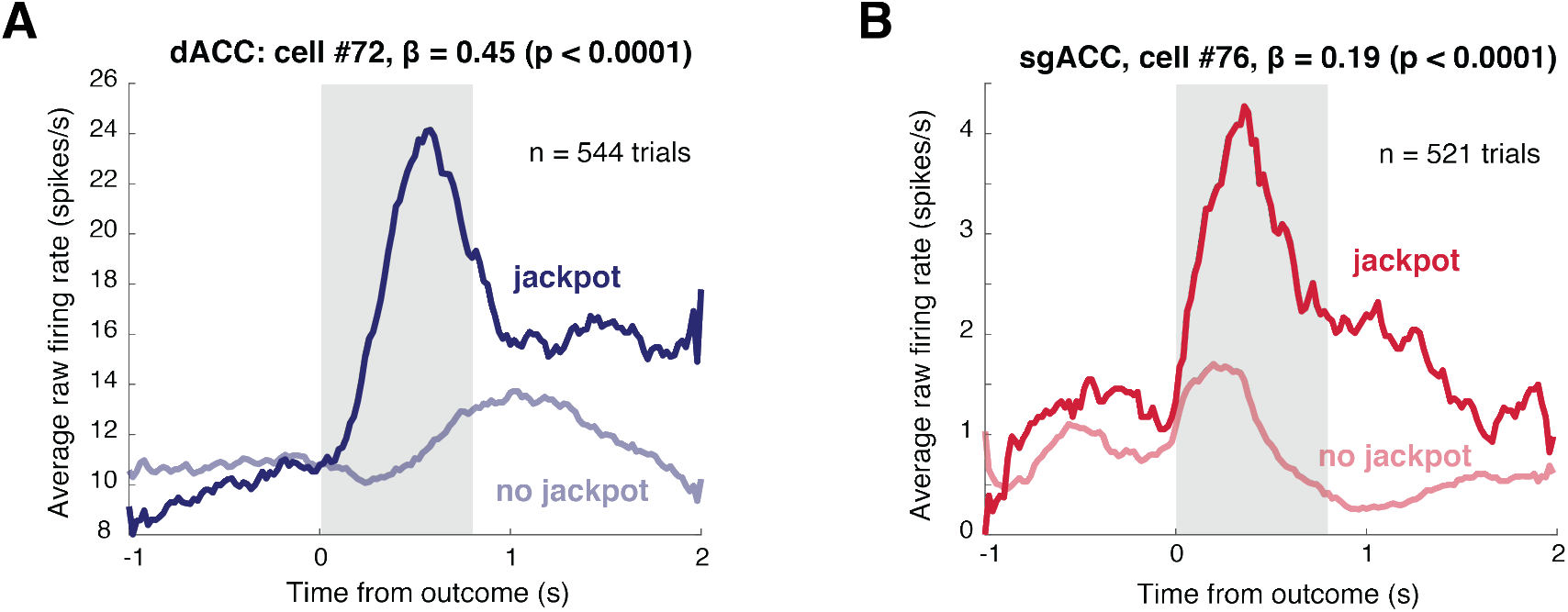
Neural responses to the anticipation of a large liquid reward (jackpot). **A:** Responses of dACC neuron with increasing firing rate before jackpot rewards. **B:** Responses of sgACC neuron with increasing firing rate before jackpot rewards. There was a bias towards this response profile in the sgACC population (see text and Table 2).

### Neurons in both regions carry spatial signals

We previously showed coding for two task variables, spatial position of offer and chosen option, in both dACC and sgACC (Strait et al., 2016). The sgACC data used in that study (n = 112 neurons) were augmented with additional neurons collected in the same animals in the same task (n = 34 additional neurons; total of 146 neurons). We still find these effects in the larger dataset. (Note that for consistency with the other analyses in this manuscript, we used a *multiple* linear regression analysis, see Methods). Specifically, sgACC encoded the position of the offer during the first offer epoch (n = 14/146 neurons, P = 0.016; two-sided binomial test), and the position of the chosen option during the post-choice epoch (n = 20/146 neurons, P < 0.0001). Results remained qualitatively similar in dACC with our new analyses. We see a significant proportion of neurons tuned to the side of the first offer in the first epoch (n = 20/129 neurons, P < 0.0001), and to the side of the chosen offer in the post-choice epoch (n = 30/129 neurons, P < 0.0001).

### No significant difference in latency of encoding chosen side between regions

We investigated latency differences between dACC and sgACC. We were particularly interested in encoding of the chosen side, since this variable defined the action ultimately taken to indicate the subject’s decision. Conventional latency analyses may be confounded due to the difference in signal-to-noise ratio between regions. This problem is especially acute when these differences are large, as they are in our sgACC and dACC datasets. In our data, signals are likely to achieve significance sooner in dACC, simply because they are stronger than in sgACC.

To avoid this confound, we developed a new procedure to control for signal-to-noise differences. We added Poisson distributed noise to our dACC dataset (mean = mean firing rate of each cell), and tested the proportion of variance explained in this new whitened dataset. If signal-to-noise in the reward epoch (when both regions were most responsive) was within 1.5% of that in sgACC, we included this dataset in our analyses. We generated 500 datasets in this manner, then performed a latency analysis the same as in Strait et al. (2015), with the slight modification that we used a multiple regression model instead of a t-test to determine when neurons first responded significantly to chosen side (this analysis takes into account neuronal responses to other variables, and is therefore more robust). We used this manipulation to account for the variance explained by other task-variables (listed in Methods). Even using this more sensitive measure and better-controlled analysis, we failed to see a significant difference between latencies across regions (t-test across regions significant for 1/500 datasets, P = 0.998).

## DISCUSSION

We examined responses of neurons in two cingulate regions, dACC and sgACC, in a token gambling task. Both areas encoded several task variables, including values of attended and remembered offers, outcomes, current token number, and, confirming an earlier study, position of offered and chosen options. Neuronal response latencies were similar in both areas. We observed two major differences between the regions: first, the task-related signal-to-noise was consistently greater in dACC than in sgACC. Second, we observed some differences in the average direction of value tuning: sgACC shows a negative bias to outcomes while dACC shows a positive one, and, prior to jackpots, sgACC shows a positive bias in activation, while dACC shows no consistent pattern. Overall, these results point to a broad functional overlap between dACC and sgACC, with differences more in emphasis than in major functional role.

Studies of functional neuroanatomy often focus on identifying each area’s unique specialization, but there are other important questions about brain regions—such as: which regions subserve a given brain function? One mental operation may be implemented by similar computations occurring simultaneously in multiple brain areas (Behrmann & Plaut, 2013; Cisek & Kalaska, 2010; Cisek, 2012; Farah, 1994; McClelland & Rumelhart, 1986). Our data suggest that risky economic choice may be mediated by computations occurring in both dACC and sgACC, and, given our earlier data, across vmPFC and VS as well (Strait et al., 2014; Strait et al., 2015). While these regions may, in principle, compute similar variables for distinct purposes, it is more parsimonious to assume, without strong evidence to the contrary, that they play similar roles.

By way of analogy, an earlier generation of scholars proposed that early and mid-level visual areas are specialized for specific aspects of form (Zeki, 1978), and that V4 is specialized for color (Essen & Zeki, 1978; Zeki, 1977; Zeki, 1980). Subsequent work disproved this idea and demonstrated that its role in color processing is not very different from those of adjacent areas (Schein et al., 1982; Motter, 1994; Van Essen & Maunsell, 1983). Its properties are similar to (but slightly more complex than) its afferent V2 and similar to (but slightly less complex than) its efferent PIT (Desimone & Schein, 1987; Desimone et al., 1985; Desimone & Duncan, 1995; Hayden & Gallant, 2013; Mirabella et al., 2007). It would be a mistake to look for color vision in any one brain region: it depends on the distributed and coordinated action of several brain regions, all of which perform many other computations unrelated to color (Desimone & Duncan, 1995; Lennie, 1998). We suspect that the same principles apply to economic choice; it reflects qualitatively similar, although quantitatively different, repeated simple computations occurring in parallel across multiple prefrontal and striatal regions (and possibly others as well). These results thus argue against a strong functional dichotomy between the two regions. For example, Bush et al. (2000) argue that the dorsal cognitive region is functionally distinct from the subgenual affective region and they may even inhibit each other. We see no evidence for a cognitive/limbic split, nor of mutual inhibition between these two regions. Clearly, other studies, using other methods, do see evidence for such a split; further work will be required to reconcile these contradictions.

At the same time, the differences we observe between dACC and sgACC argue against a mass action view of brain function. The differences we see are consistent with results of both neuroimaging and lesion studies supporting some functional difference between the areas, and with the neuroanatomy, which shows different patterns of connectivity (Heilbronner & Haber, 2014 Vogt et al., 1995). The difference in signal-to-noise is particularly intriguing. It may reflect a greater selectivity for choice tasks in the dorsal region; presumably there are other tasks that would drive sgACC more effectively than dACC (however, we would expect these differences to be quantitative, not qualitative). Another possibility is that economic decision-making, while distributed, has a fan-in structure and that dACC, which is presumably later in the hierarchy, has a more concentrated and thus stronger set of task signals.

Our findings generally confirm and extend the existing literature on sgACC. Monosov & Hikosaka (2012), like us, find a bias towards encoding of negative outcomes in area 25. We show that this finding extends to secondary (token) reinforcers. They report no encoding of probability, and thus of integrated value; our finding of integrated value encoding thus suggests that sgACC may play a more direct economic role, especially in decision-making tasks. Amemori & Graybiel (2012) recorded in pregenual cingulum, in a region that is rostral to our region, but has a small overlap (see Methods). They found a subzone, which may overlap with our subgenual recording site, biased towards negative encodings. Our high fraction of subgenual neurons encoding upcoming large rewards (jackpots) provide confirmation for the idea, proposed by Rudebeck and colleagues (2014), that sgACC serves to sustain autonomic arousal in anticipation of a reward. The phasic, task-relevant signals in sgACC, and its direct role in reward-processing and anticipation, suggest that the role for subgenual neurons extends beyond the control of basic arousal functions like sleep (Rolls et al., 2003). Our findings also cohere with the findings relating activity in sgACC to negative affect and depression. Overactivity in this region correlates with depressive symptoms (Drevets, 2002; Mayberg et al., 2000) and with transient sadness in healthy subjects (Mayberg et al., 1999); chronic stimulation of sgACC can alleviate the symptoms of depression (Mayberg et al., 2005). Negative mood inducing stimuli tend to activate sgACC (George et al., 1995; Mayberg et al., 1997; Mayberg et al., 1999), and structural abnormalities in sgACC correlate with mood disorders (Botteron et al., 2002; Coryell et al., 2005; Drevets et al., 1997).

These findings also have implications for our understanding of dACC as well. The role of dACC in economic choice remains disputed. Some research indicates that it serves as the site of choice (Rangel & Hare, 2010); our work confirms this idea but, in the broader context of data from other studies, suggests it does not play a unique role. Other research suggests that dACC plays a specifically post-decisional role, in part because it preferentially encodes the value of unchosen options (Blanchard & Hayden, 2014; Boorman et al., 2011; Rushworth et al., 2012; Hayden et al., 2009). Our results here suggest that dACC may encode predecisional variables: the values of both chosen and unchosen options – at least before the decision is made. Other work links dACC to persistence – that is, in maintenance of value encoding until the time of reward to allow stable behavior (Hillman & Bilkey, 2013, Chudasama et al., 2013; Picton et al., 2007; Blanchard et al., 2015; Shidara and Richmond, 2002). While we did not use a persistence task, our results do endorse the idea that dACC serves to encode prospective rewards, suggesting that it does serve this function although not necessarily purely for the purpose of persistence.

Overall, these results invite speculation on whether one can ascribe a single integrated role for the cingulum (Hayden & Platt, 2009). On one hand, the broad overlap between dACC and sgACC that we see suggests it may have a general economic role. On the other, our findings in a similar task in vmPFC and VS suggest this role may not be unique to the cingulate cortex (Strait et al., 2014; Strait et al., 2015). Our results also have some similarity to findings from the posterior cingulate cortex. Neural responses in that region are associated with several economic variables, including offer and outcome variables, as well as learning and control variables (Dean & Platt, 2003; Hayden et al., 2009; Heilbronner & Platt, 2013; McCoy et al., 2003; Heilbronner et al., 2011). Neurons in sgACC tend to have poorer signal-to-noise than dACC, much like neurons in PCC (Hayden et al., 2008), suggesting they may have some affinity. Still, we consider the question of whether there is a general and unique function for the cingulum to be unanswered.

## ACKNOWLEDGEMENTS

We thank Caleb Strait for assistance designing the task, Tommy Blanchard, and Alex Thomé for useful discussions, and Meghan Castagno, Giuliana Loconte and Marc Mancarella for assistance in data collection. This research was supported by a NIH R01 (DA037229) and a NSF CAREER award to BYH.

## MATERIALS AND METHODS

Some of the data for both dACC and sgACC recordings were published (Strait et al., 2016); all analyses presented here are new.

### Surgical Procedures

All procedures were approved by the University Committee on Animal Resources at the University of Rochester and were designed and conducted in compliance with the Public Health Service’s Guide for the Care and Use of Animals. Two male rhesus macaques (*Macaca mulatta*: subject B age 5y. 7mo.; subject J age 6y. 7mo.) served as subjects. A small prosthesis for holding the head was used. Animals were habituated to laboratory conditions and then trained to perform oculomotor tasks for liquid reward. A Cilux recording chamber (Crist Instruments) was placed over the dACC. Position was verified by magnetic resonance imaging with the aid of a Brainsight system (Rogue Research Inc.). Animals received appropriate analgesics and antibiotics after all procedures. Throughout both behavioral and physiological recording sessions, the chamber was kept sterile with regular antibiotic washes and sealed with sterile caps. All recordings were performed during the animals’ light cycle between 8am and 5pm.

### Recording Site

We approached dACC through a standard recording grid (Crist Instruments). We defined dACC according to the Paxinos atlas (Paxinos et al., 2000). Roughly, we recorded from a ROI lying within the coronal planes situated between 29.50 and 34.50 mm rostral to interaural plane, the horizontal planes situated between 4.12 to 7.52 mm from the brain’s dorsal surface, and the sagittal planes between 0 and 5.24 mm from medial wall. Our recordings were made from a central region within this zone. We confirmed recording location before each recording session using our Brainsight system with structural magnetic resonance images taken before the experiment. Neuroimaging was performed at the Rochester Center for Brain Imaging, on a Siemens 3T MAGNETOM Trio Tim using 0.5 mm voxels. We confirmed recording locations by listening for characteristic sounds of white and gray matter during recording, which in all cases matched the loci indicated by the Brainsight system.

We approached sgACC using similar equipment, in a similar manner. We defined sgACC as lying within the coronal planes situated between 24 and 36 mm rostral to interaural plane, the horizontal planes situated between 17.33 to 25.12 mm from the brain’s dorsal surface, and the sagittal planes between 0 and 5.38 mm from the medial wall. Our recordings were made from a central region within this zone. We again confirm recording locations using structural magnetic resonance images, Brainsight, and listening to characteristic sounds of white and gray matter. These regions correspond to area 25 as identified by Paxinos (n = 118 neurons) and also to the most caudal portion of area 32 (n = 28 neurons).

### Electrophysiological Techniques

Single electrodes (Frederick Haer & Co., impedance range 0.8 to 4 MU) were lowered using a microdrive (NAN Instruments) until waveforms of between one and three neuron(s) were isolated. Individual action potentials were isolated on a Plexon system (Plexon, Inc.). Neurons were selected for study solely on the basis of the quality of isolation; we never pre-selected based on task-related response properties. All collected neurons for which we managed to obtain at least 250 trials were analyzed.

### Eye Tracking and Reward Delivery

Eye position was sampled at 1,000 Hz by an infrared eye-monitoring camera system (SR Research). Stimuli were controlled by a computer running Matlab (Mathworks) with Psychtoolbox (Brainard,1997) and Eyelink Toolbox (Cornelissen et al., 2002). Visual stimuli were colored rectangles on a computer monitor placed 57 cm from the animal and centered on its eyes (Figure 1A). A standard solenoid valve controlled the duration of juice delivery. The relationship between solenoid open time and juice volume was established and confirmed before, during, and after recording.

### Behavioral Task

Monkeys performed a two-option gambling task. The task was similar to one we have used previously (Strait et al., 2014; Strait et al., 2015), with two major differences: (1) monkeys gambled for virtual tokens—rather than liquid—rewards, and (2) outcomes could be losses as well as wins.

Two offers were presented on each trial. Each offer was represented by a rectangle 300 pixels tall and 80 pixels wide (11.35° of visual angle tall and 4.08° of visual angle wide). 20% of options were safe (100% probability of either 0 or 1 token), while the remaining 80% were gambles. Safe offers were entirely red (0 tokens) or blue (1 token). The size of each portion indicated the probability of the respective reward. Each gamble rectangle was divided horizontally into a top and bottom portion, each colored according to the token reward offered. Gamble offers were thus defined by three parameters: two possible token outcomes, and probability of the top outcome (the probability of the bottom was strictly determined by the probability of the top). The top outcome was 10%, 30%, 50%, 70% or 90% likely.

Six initially unfilled circles arranged horizontally at the bottom of the screen indicated the number of tokens to be collected before the subject obtained a liquid reward. These circles were filled appropriately at the end of each trial, according to the outcome of that trial. When 6 or more tokens were collected, the tokens were covered with a solid rectangle while a liquid reward was delivered. Tokens beyond 6 did not carry over, nor could number of tokens fall below zero.

On each trial, one offer appeared on the left side of the screen and the other appeared on the right. Offers were separated from the fixation point by 550 pixels (27.53° of visual angle). The side of the first offer (left and right) was randomized by trial. Each offer appeared for 600 ms and was followed by a 150 ms blank period. Monkeys were free to fixate upon the offers when they appeared (and in our observations almost always did so). After the offers were presented separately, a central fixation spot appeared and the monkey fixated on it for 100 ms. Following this, both offers appeared simultaneously and the animal indicated its choice by shifting gaze to its preferred offer and maintaining fixation on it for 200 ms. Failure to maintain gaze for 200 ms did not lead to the end of the trial, but instead returned the monkey to a choice state; thus, monkeys were free to change their mind if they did so within 200 ms (although in our observations, they seldom did so). A successful 200 ms fixation was followed by a 750 ms delay, after which the gamble was resolved and a small reward (100 µL) was delivered—regardless of the outcome of the gamble—to sustain motivation. This small reward was delivered within a 300 ms window. If 6 tokens were collected, a delay of 500 ms was followed by a large liquid reward (300 µL) within a 300 ms window, followed by a random inter-trial interval (ITI) between 0.5 and 1.5 s. If 6 tokens were not collected, subjects proceeded immediately to the ITI.

Each gamble included at least one positive or zero-outcome, ensuring that every gamble carried the possibility of a win. This decreased the number of trivial choices presented to subjects, and maintained motivation.

### Statistical Methods

**PSTHs** were constructed by aligning spike rasters to the task event of interest (offer 1 appearance, choice or feedback) and averaging firing rates across multiple trials. Firing rates were calculated in 20 ms bins but were generally analyzed in longer (500-1000 ms) epochs. Plots were generated in Matlab, and use the Matlab function *smooth* with a smoothing factor of 20.

Firing rates were **normalized** by subtracting the mean and dividing by the standard deviation of the entire neuron’s psth.

We test for significant tuning and assess variance explained using a **multiple linear regression**, including the following task-relevant variables: expected value of offers 1 and 2, the number of tokens collected as of the beginning of the trial, the side the first offer appeared on, conflict between offer values (defined as the absolute difference between them), the side of the chosen offer, the outcome of the trial (in tokens), and the probability of that outcome (a measure of surprise).

**Analysis epochs** were chosen *a priori*, before data analysis began, to reduce the likelihood of p-hacking. The first and second offer epochs were defined as the 500 ms epoch beginning 100 ms after the offer was presented, to account for information processing time. The choice epoch was the 1-second epoch before choice was indicated using an express saccade. The reward epoch was defined as the 800 ms epoch following the resolution of a gamble: this was when feedback and a small liquid reward were given (regardless of trial outcome), followed by a 500 ms delay (this was part or all of the intertrial interval on non-jackpot trials, and a pre-jackpot delay on jackpot trials).

All **fractions of tuned neurons** were tested for significance using a two-sided binomial test. All binomial tests throughout the manuscript were two-sided.

All **sliding window plots** use a sliding 500 ms window, computing the fraction of tuned neurons in the population every 20 ms. Points on the plot are aligned to the start of this window. Significance is assessed using a multiple linear regression including all task-variables (mentioned above). Plots were smoothed using the Matlab function *smooth* and a smoothing factor of 20.

We use **beta correlation analyses using stepwise regression** to assess whether neurons represent two variables (or the same variable at different time periods) using similar / orthogonal / opposing formats, in overlapping / orthogonal / distinct populations. In these analyses, we first find the regression coefficients associated with the variables in question, then find the Pearson correlation coefficient between these.

We noticed that, depending on which variables we included in the regression, the results of our beta correlation analyses sometimes differed. We therefore used a stepwise regression model as an objective method of determining which variables should be included in the final regression for each individual neuron. We first include all relevant task variables, including the variables of interest, in this stepwise regression. We then perform a (non-stepwise) multiple linear regression analysis again using only the variables obtained from the previous step, as well as the variables of interest. We do this to obtain a regression coefficient corresponding to the variable of interest for each neuron, regardless of whether the coefficient for this variable achieved significance in the previous step.

The Pearson correlation coefficient between signed regression coefficients indicate whether variables were represented in a similar *format* i.e. directionality of tuning across the population. A positive correlation indicates a preservation of directionality, while a negative correlation indicates variables were represented in opposing directionality of firing rate modulation. No correlation suggests orthogonal formats, but we draw no strong conclusions from these.

Similarly, the Pearson correlation coefficient between unsigned regression coefficients indicates whether similar neuronal populations tended to be involved in encoding the two variables in question. A positive correlation indicates overlapping populations, while a negative correlation indicates separate ones. A lack of correlation suggests orthogonal populations (i.e. encoding one variable does not affect a neuron’s likelihood of encoding the other variable), but we again draw no strong conclusions from this null result.

We performed a **cross-validation procedure** similar to that in Blanchard et al. (Blanchard et al., 2015) on our sgACC dataset to determine whether our signal in the first and second epochs was strong enough to detect correlations. We split trials into two random sets (even and odd), and computed the regression coefficients for each individual neuron in response to offer 1 value in both epochs 1 and 2. We then obtained the Pearson correlation coefficient of these signed regression coefficients within each epoch. We see no correlation in the epoch we analyzed for offer 1 memory signals (i.e. epoch 2: r = 0.16, P = 0.058, Pearson correlation coefficient). The failure of this cross-validation procedure to detect a significant correlation in sgACC indicates that we did not have sufficient signal in our dataset to detect correlation effects, even if they existed. This procedure prevents us from drawing strong conclusions regarding the *absence* of a signal in this region.

## REFERENCES

Akaishi, R., & Hayden, B. Y. (2016). A Spotlight on Reward. Neuron, 90(6), 1148–1150. http://doi.org/10.1016/j.neuron.2016.06.008

Allman, J. M., Hakeem, A., Erwin, J. M., Nimchinsky, E., & Hof, P. (2001). The Anterior Cingulate Cortex. Annals of the New York Academy of Sciences, 935(1), 107–117. http://doi.org/10.1111/j.1749-6632.2001.tb03476.x

Amemori, K. I., & Graybiel, A. M. (2012). Localized microstimulation of primate pregenual cingulate cortex induces negative decision-making. Nature neuroscience, 15(5), 776–785.

Amiez, C., Joseph, J.-P., & Procyk, E. (2005). Anterior cingulate error-related activity is modulated by predicted reward. European Journal of Neuroscience, 21(12), 3447–3452. http://doi.org/10.1111/j.1460-9568.2005.04170.x

Behrmann, M., & Plaut, D. C. (2013). Distributed circuits, not circumscribed centers, mediate visual recognition. Trends in Cognitive Sciences, 17(5), 210–219. http://doi.org/10.1016/j.tics.2013.03.007

Blanchard, T. C., & Hayden, B. Y. (2014). Neurons in Dorsal Anterior Cingulate Cortex Signal Postdecisional Variables in a Foraging Task. Journal of Neuroscience, 34(2), 646–655. http://doi.org/10.1523/JNEUROSCI.3151-13.2014

Blanchard, T. C., Hayden, B. Y., & Bromberg-Martin, E. S. (2015). Orbitofrontal Cortex Uses Distinct Codes for Different Choice Attributes in Decisions Motivated by Curiosity. Neuron, 85(3), 602–614. http://doi.org/10.1016/j.neuron.2014.12.050

Blanchard, T. C., Strait, C. E., & Hayden, B. Y. (2015). Ramping ensemble activity in dorsal anterior cingulate neurons during persistent commitment to a decision. Journal of Neurophysiology, 114(4), 2439–2449. http://doi.org/10.1152/jn.00711.2015

Blumberg, H. P., Stern, E., Martinez, D., Ricketts, S., De Asis, J., White, T., … & Silbersweig, D. A. (2000). Increased anterior cingulate and caudate activity in bipolar mania. Biological psychiatry, 48(11), 1045–1052.

Boorman ED, Behrens TE, Rushworth MF (2011) Counterfactual choice and learning in a neural network centered on human lateral frontopolar cortex. PLoS Biol 9:e1001093. CrossRef Medline

Botteron, K. N., Raichle, M. E., Drevets, W. C., Heath, A. C., & Todd, R. D. (2002). Volumetric reduction in left subgenual prefrontal cortex in early onset depression. Biological Psychiatry, 51(4), 342–344. http://doi.org/10.1016/S0006-3223(01)01280-X

Bouras, C., Kövari, E., Hof, P. R., Riederer, B. M., & Giannakopoulos, P. (2001). Anterior cingulate cortex pathology in schizophrenia and bipolar disorder. Acta Neuropathologica, 102(4), 373–379. http://doi.org/10.1007/s004010100392

Bush, G., Frazier, J. A., Rauch, S. L., Seidman, L. J., Whalen, P. J., Jenike, M. A., et al. (1999). Anterior cingulate cortex dysfunction in attention-deficit/hyperactivity disorder revealed by fMRI and the counting stroop. Biological Psychiatry, 45(12), 1542–1552. http://doi.org/10.1016/S0006-3223(99)00083-9

Bush, G., Luu, P., & Posner, M. I. (2000). Cognitive and emotional influences in anterior cingulate cortex. Trends in Cognitive Sciences, 4(6), 215–222. http://doi.org/10.1016/S1364-6613(00)01483-2

Bush, G., Vogt, B. A., Holmes, J., Dale, A. M., Greve, D., Jenike, M. A., & Rosen, B. R. (2002). Dorsal anterior cingulate cortex: a role in reward-based decision making. Proceedings of the National Academy of Sciences, 99(1), 523–528. http://doi.org/10.1073/pnas.012470999

Carter, C. S., & van Veen, V. (2007). Anterior cingulate cortex and conflict detection: An update of theory and data. Cognitive, Affective, & Behavioral Neuroscience, 7(4), 367–379. http://doi.org/10.3758/CABN.7.4.367

Chudasama, Y., Daniels, T. E., Gorrin, D. P., Rhodes, S. E., Rudebeck, P. H., & Murray, E. A. (2013). The role of the anterior cingulate cortex in choices based on reward value and reward contingency. Cerebral Cortex, 23(12), 2884–2898.

Cisek, P. (2012). Making decisions through a distributed consensus. Current Opinion in Neurobiology, 22(6), 927–936. http://doi.org/10.1016/j.conb.2012.05.007

Cisek, P., & Kalaska, J. F. (2010). Neural Mechanisms for Interacting with a World Full of Action Choices. Dx.Doi.org, 33(1), 269–298. http://doi.org/10.1146/annurev.neuro.051508.135409

Coryell, W., Nopoulos, P., Drevets, W., Wilson, T., & Andreasen, N. C. (2005). Subgenual Prefrontal Cortex Volumes in Major Depressive Disorder and Schizophrenia: Diagnostic Specificity and Prognostic Implications. American Journal of Psychiatry, 162(9), 1706–1712. http://doi.org/10.1176/appi.ajp.162.9.1706

Cotter, D., Mackay, D., Landau, S., Kerwin, R., & Everall, I. (2001). Reduced Glial Cell Density and Neuronal Size in the Anterior Cingulate Cortex in Major Depressive Disorder. Archives of General Psychiatry, 58(6), 545–553. http://doi.org/10.1001/archpsyc.58.6.545

Dean, H. L., & Platt, M. L. (2003). Spatial representations in posterior cingulate cortex. Journal of Vision, 3(9), 427–427. http://doi.org/10.1167/3.9.427

Derbyshire, S. W. G., Vogt, B. A., & Jones, A. K. P. (1998). Pain and Stroop interference tasks activate separate processing modules in anterior cingulate cortex. Experimental Brain Research, 118(1), 52–60. http://doi.org/10.1007/s002210050254

Desimone, R., & Duncan, J. (1995). Neural mechanisms of selective visual attention. Annual Review of Neuroscience.

Desimone, R., & Schein, S. J. (1987). Visual properties of neurons in area V4 of the macaque: sensitivity to stimulus form. Journal of Neurophysiology, 57(3), 835–868.

Desimone, R., Schein, S. J., Moran, J., & Ungerleider, L. G. (1985). Contour, color and shape analysis beyond the striate cortex. Vision Research, 25(3), 441–452. http://doi.org/10.1016/0042-6989(85)90069-0

Drevets, W. (2002). Functional anatomical correlates of antidepressant drug treatment assessed using PET measures of regional glucose metabolism. European Neuropsychopharmacology, 12(6), 527–544. http://doi.org/10.1016/S0924-977X(02)00102-5

Drevets, W. C., Price, J. L., Simpson, J. R., Todd, R. D., Reich, T., Vannier, M., & Raichle, M. E. (1997). Subgenual prefrontal cortex abnormalities in mood disorders.

Drevets, W. C., Savitz, J., & Trimble, M. (2008). The subgenual anterior cingulate cortex in mood disorders. CNS Spectrums, 13(8), 663–681.

Dum, R. P., Levinthal, D. J., & Strick, P. L. (2016). Motor, cognitive, and affective areas of the cerebral cortex influence the adrenal medulla. Proceedings of the National Academy of Sciences of the United States of America, 113(35), 9922–9927. http://doi.org/10.1073/pnas.1605044113

Essen, D. C., & Zeki, S. M. (1978). The topographic organization of rhesus monkey prestriate cortex. The Journal of Physiology, 277, 193–226. http://doi.org/10.1111/(ISSN)1469-7793

Farah, M. J. (1994). Neuropsychological inference with an interactive brain: A critique of the “locality” assumption. Behavioral and Brain Sciences, 17(1), 43–61. http://doi.org/10.1017/S0140525X00033306

Forman, S. D., Dougherty, G. G., Casey, B. J., Siegle, G. J., Braver, T. S., Barch, D. M., … & Lorensen, E. (2004). Opiate addicts lack error-dependent activation of rostral anterior cingulate. Biological psychiatry, 55(5), 531–537.

Freedman, L. J., Insel, T. R., & Smith, Y. (2000). Subcortical projections of area 25 (subgenual cortex) of the macaque monkey. Journal of Comparative Neurology, 421(2), 172–188. http://doi.org/10.1002/(SICI)1096-9861(20000529)421:2<172::AID-CNE4>3.0.CO;2-8

George, M. S., Ketter, T. A., Parekh, P. I., Horwitz, B., Herscovitch, P., & Post, R. M. (2006). Brain activity during transient sadness and happiness in healthy women. American Journal of Psychiatry, 152(3), 341–351. http://doi.org/10.1176/ajp.152.3.341

Goldstein, R. Z., Tomasi, D., Rajaram, S., Cottone, L. A., Zhang, L., Maloney, T., et al. (2007). Role of the anterior cingulate and medial orbitofrontal cortex in processing drug cues in cocaine addiction. Neuroscience, 144(4), 1153–1159. http://doi.org/10.1016/j.neuroscience.2006.11.024

Graybiel, A. M., & Rauch, S. L. (2000). Toward a Neurobiology of Obsessive-Compulsive Disorder. Neuron, 28(2), 343–347. http://doi.org/10.1016/S0896-6273(00)00113-6

Hayden, B., & Gallant, J. (2013). Working Memory and Decision Processes in Visual Area V4. Frontiers in Neuroscience, 7. http://doi.org/10.3389/fnins.2013.00018

Hayden, B. Y., Heilbronner, S. R., Nair, A. C., & Platt, M. L. (2008). Cognitive influences on risk-seeking by rhesus macaques. Judgment and Decision Making, 3(5), 389–395.

Hayden, B. Y., Nair, A. C., McCoy, A. N., & Platt, M. L. (2008). Posterior Cingulate Cortex Mediates Outcome-Contingent Allocation of Behavior. Neuron, 60(1), 19–25. http://doi.org/10.1016/j.neuron.2008.09.012

Hayden, B. Y., Pearson, J. M., & Platt, M. L. (2009). Fictive reward signals in the anterior cingulate cortex. science, 324(5929), 948–950.

Hayden, B. Y., Pearson, J. M., & Platt, M. L. (2011). Neuronal basis of sequential foraging decisions in a patchy environment. Nature neuroscience, 14(7), 933–939.

Hayden, B. Y., & Platt, M. L. (2007). Temporal Discounting Predicts Risk Sensitivity in Rhesus Macaques. Current Biology, 17(1), 49–53. http://doi.org/10.1016/j.cub.2006.10.055

Hayden, B. Y., & Platt, M. L. (2009). Cingulate cortex. Encyclopaedia of Neuroscience. http://doi.org/10.3389/fnhum.2012.00124/abstract

Hayden, B. Y., Smith, D. V., & Platt, M. L. (2009). Electrophysiological correlates of default-mode processing in macaque posterior cingulate cortex. Proceedings of the National Academy of Sciences of the United States of America, 106(14), 5948–5953. http://doi.org/10.1073/pnas.0812035106

Heilbronner, S. R., & Haber, S. N. (2014). Frontal cortical and subcortical projections provide a basis for segmenting the cingulum bundle: implications for neuroimaging and psychiatric disorders. The Journal of Neuroscience, 34(30), 10041–10054.

Heilbronner, S. R., & Hayden, B. (2015). Contextual factors explain risk-seeking preferences in rhesus monkeys. Decision Making under Uncertainty, 91.

Heilbronner, S. R., & Hayden, B. Y. (2016). Dorsal Anterior Cingulate Cortex: A Bottom-Up View. Annual Review of Neuroscience.

Heilbronner, S. R., & Platt, M. L. (2013). Causal Evidence of Performance Monitoring by Neurons in Posterior Cingulate Cortex during Learning. Neuron, 80(6), 1384–1391. http://doi.org/10.1016/j.neuron.2013.09.028

Hillman, K. L., & Bilkey, D. K. (2013). Persisting through subjective effort: A key role for the anterior cingulate cortex? Behavioral and Brain Sciences, 36(6), 691–692. http://doi.org/10.1017/S0140525X13001039

Hosokawa, T., Kennerley, S. W., Sloan, J., & Wallis, J. D. (2013). Single-neuron mechanisms underlying cost-benefit analysis in frontal cortex. The Journal of Neuroscience, 33(44), 17385–17397. http://doi.org/10.1523/JNEUROSCI.2221-13.2013

Hunt, L. T., Behrens, T. E., Hosokawa, T., Wallis, J. D., & Kennerley, S. W. (2015). Capturing the temporal evolution of choice across prefrontal cortex. Elife, 4, e11945.

Johansen-Berg, H., Gutman, D. A., Behrens, T. E. J., Matthews, P. M., Rushworth, M. F. S., Katz, E., et al. (2008). Anatomical Connectivity of the Subgenual Cingulate Region Targeted with Deep Brain Stimulation for Treatment-Resistant Depression. Cerebral Cortex, 18(6), 1374–1383. http://doi.org/10.1093/cercor/bhm167

Kennerley, S. W., Walton, M. E., Behrens, T. E. J., Buckley, M. J., & Rushworth, M. F. S. (2006). Optimal decision making and the anterior cingulate cortex. Nature Neuroscience, 9(7), 940–947. http://doi.org/10.1038/nn1724

Kerns, J. G., Cohen, J. D., MacDonald, A. W., Cho, R. Y., Stenger, V. A., & Carter, C. S. (2004). Anterior Cingulate Conflict Monitoring and Adjustments in Control. Science, 303(5660), 1023–1026. http://doi.org/10.1126/science.1089910

Lak, A., Stauffer, W. R., & Schultz, W. (2014). Dopamine prediction error responses integrate subjective value from different reward dimensions. Proceedings of the National Academy of Sciences of the United States of America, 111(6), 2343–2348. http://doi.org/10.1073/pnas.1321596111

Lennie, P. (1998). Single Units and Visual Cortical Organization. Perception, 27(8), 889–935. http://doi.org/10.1068/p270889

Mayberg, H. S., Brannan, S. K., Mahurin, R. K., Jerabek, P. A., Brickman, J. S., Tekell, J. L., et al. (1997). Cingulate function in depression: a potential predictor of treatment response. NeuroReport, 8(4), 1057.

Mayberg, H. S., Brannan, S. K., Tekell, J. L., Silva, J. A., Mahurin, R. K., McGinnis, S., & Jerabek, P. A. (2000). Regional metabolic effects of fluoxetine in major depression: serial changes and relationship to clinical response. Biological Psychiatry, 48(8), 830–843. http://doi.org/10.1016/S0006-3223(00)01036-2

Mayberg, H. S., Liotti, M., & Brannan, S. K. (1999). Reciprocal limbic-cortical function and negative mood: converging PET findings in depression and normal sadness. American Journal of ….

Mayberg, H. S., Lozano, A. M., Voon, V., McNeely, H. E., Seminowicz, D., Hamani, C., et al. (2005). Deep Brain Stimulation for Treatment-Resistant Depression. Neuron, 45(5), 651–660. http://doi.org/10.1016/j.neuron.2005.02.014

McClelland, J. L., Rumelhart, D. E., & Hinton, G. E. (1986). The appeal of parallel distributed processing. MIT Press, Cambridge MA, 3–44.

McCoy, A. N., Crowley, J. C., Haghighian, G., Dean, H. L., & Platt, M. L. (2003). Saccade Reward Signals in Posterior Cingulate Cortex. Neuron, 40(5), 1031–1040. http://doi.org/10.1016/S0896-6273(03)00719-0

McCoy, A. N., & Platt, M. L. (2005). Risk-sensitive neurons in macaque posterior cingulate cortex. Nature neuroscience, 8(9), 1220–1227.

Mirabella, G., Bertini, G., Samengo, I., Kilavik, B. E., Frilli, D., Libera, Della, C., & Chelazzi, L. (2007). Neurons in Area V4 of the Macaque Translate Attended Visual Features into Behaviorally Relevant Categories. Neuron, 54(2), 303–318. http://doi.org/10.1016/j.neuron.2007.04.007

Monosov, I. E., & Hikosaka, O. (2012). Regionally distinct processing of rewards and punishments by the primate ventromedial prefrontal cortex. The Journal of Neuroscience, 32(30), 10318–10330. http://doi.org/10.1523/JNEUROSCI.1801-12.2012

Motter, B. C. (1994). Neural correlates of attentive selection for color or luminance in extrastriate area V4. Journal of Neuroscience **14,** 2178–2189.

Neubert, F.-X., Mars, R. B., Sallet, J., & Rushworth, M. F. S. (2015). Connectivity reveals relationship of brain areas for reward-guided learning and decision making in human and monkey frontal cortex. Proceedings of the National Academy of Sciences, 112(20), E2695–E2704. http://doi.org/10.1073/pnas.1410767112

Öngür, D., Drevets, W. C., & Price, J. L. (1998). Glial reduction in the subgenual prefrontal cortex in mood disorders. Proceedings of the National Academy of Sciences, 95(22), 13290–13295. http://doi.org/10.1073/pnas.95.22.13290

Paus, T. (2001). Primate Anterior Cingulate Cortex: Where Motor Control, Drive and Cognition Interface. Nature Reviews Neuroscience, 1–8.

Picton, T. W., Stuss, D. T., Alexander, M. P., Shallice, T., Binns, M. A., & Gillingham, S. (2007). Effects of focal frontal lesions on response inhibition. Cerebral Cortex, 17(4), 826–838.

Procyk, E., Tanaka, Y. L., & Joseph, J. P. (2000). Anterior cingulate activity during routine and non-routine sequential behaviors in macaques. Nature Neuroscience, 3(5), 502–508. http://doi.org/10.1038/74880

Quilodran, R., Rothé, M., & Procyk, E. (2008). Behavioral Shifts and Action Valuation in the Anterior Cingulate Cortex. Neuron, 57(2), 314–325. http://doi.org/10.1016/j.neuron.2007.11.031

Rangel, A., & Hare, T. (2010). Neural computations associated with goal-directed choice. Current Opinion in Neurobiology, 20(2), 262–270. http://doi.org/10.1016/j.conb.2010.03.001

Rich, E. L., & Wallis, J. D. (2016). Decoding subjective decisions from orbitofrontal cortex. Nature Neuroscience, 19(7), 973–980. http://doi.org/10.1038/nn.4320

Rolls, E. T., Inoue, K., & Browning, A. (2003). Activity of primate subgenual cingulate cortex neurons is related to sleep. Journal of Neurophysiology, 90(1), 134–142.

Rudebeck, P. H., Putnam, P. T., Daniels, T. E., Yang, T., Mitz, A. R., Rhodes, S. E., & Murray, E. A. (2014). A role for primate subgenual cingulate cortex in sustaining autonomic arousal. Proceedings of the National Academy of Sciences, 111(14), 5391–5396.

Rushworth, M. F. S., Hadland, K. A., Paus, T., & Sipila, P. K. (2002). Role of the Human Medial Frontal Cortex in Task Switching: A Combined fMRI and TMS Study. Journal of Neurophysiology, 87(5), 2577–2592. http://doi.org/10.1016/S0028-3932(97)00003-1

Rushworth, M. F., Kolling, N., Sallet, J., & Mars, R. B. (2012). Valuation and decision-making in frontal cortex: one or many serial or parallel systems? Current Opinion in Neurobiology, 22(6), 946–955. http://doi.org/10.1016/j.conb.2012.04.011

Schein, S. J., Marrocco, R. T., & De Monasterio, F. M. (1982). Is there a high concentration of color-selective cells in area V4 of monkey visual cortex? Journal of Neurophysiology, 47(2), 193–213. http://doi.org/10.1152/jn.00847.2015

Seo, H., & Lee, D. (2007). Temporal filtering of reward signals in the dorsal anterior cingulate cortex during a mixed-strategy game. The Journal of Neuroscience.

Shenhav, A., Botvinick, M. M., & Cohen, J. D. (2013). The Expected Value of Control: An Integrative Theory of Anterior Cingulate Cortex Function. Neuron, 79(2), 217–240. http://doi.org/10.1016/j.neuron.2013.07.007

Shidara, M., & Richmond, B. J. (2002). Anterior Cingulate: Single Neuronal Signals Related to Degree of Reward Expectancy. Science, 296(5573), 1709–1711. http://doi.org/10.1126/science.1069504

Strait, C. E., Blanchard, T. C., & Hayden, B. Y. (2014). Reward Value Comparison via Mutual Inhibition in Ventromedial Prefrontal Cortex. Neuron, 82(6), 1357–1366. http://doi.org/10.1016/j.neuron.2014.04.032

Strait, C. E., Sleezer, B. J., Blanchard, T. C., Azab, H., Castagno, M. D., & Hayden, B. Y. (2016). Neuronal selectivity for spatial positions of offers and choices in five reward regions. Journal of neurophysiology, 115(3), 1098–1111.

Strait, C. E., Sleezer, B. J., & Hayden, B. Y. (2015). Signatures of Value Comparison in Ventral Striatum Neurons. PLoS Biol, 13(6), e1002173. http://doi.org/10.1371/journal.pbio.1002173

Van Essen, D. C., & Maunsell, J. H. R. (1983). Hierarchical organization and functional streams in the visual cortex. Trends in Neurosciences, 6, 370–375. http://doi.org/10.1016/0166-2236(83)90167-4

Vogt, B. A., Finch, D. M., & Olson, C. R. (1992). Functional Heterogeneity in Cingulate Cortex: The Anterior Executive and Posterior Evaluative Regions. Cerebral Cortex, 2(6), 435–443. http://doi.org/10.1093/cercor/2.6.435-a

Vogt, B. A., Nimchinsky, E. A., Vogt, L. J., & Hof, P. R. (1995). Human cingulate cortex: Surface features, flat maps, and cytoarchitecture. Journal of Comparative Neurology, 359(3), 490–506. http://doi.org/10.1002/cne.903590310

Wallis, J. D., & Kennerley, S. W. (2010). Heterogeneous reward signals in prefrontal cortex. Current Opinion in Neurobiology, 20(2), 191–198. http://doi.org/10.1016/j.conb.2010.02.009

Yamada, H., Tymula, A., Louie, K., & Glimcher, P. W. (2013). Thirst-dependent risk preferences in monkeys identify a primitive form of wealth. Proceedings of the National Academy of Sciences of the United States of America, 110(39), 15788–15793. http://doi.org/10.1073/pnas.1308718110

Zeki, S. M. (1977). Colour Coding in the Superior Temporal Sulcus of Rhesus Monkey Visual Cortex. Proceedings of the Royal Society B: Biological Sciences, 197(1127), 195–223. http://doi.org/10.1098/rspb.1977.0065

Zeki, S. M. (1978). Functional specialisation in the visual cortex of the rhesus monkey. Nature, 274(5670), 423–428.

Zeki, S. (1980). The representation of colours. Nature, 284, 413.

